# Aerodynamic Characteristics and RNA Concentration of SARS-CoV-2 Aerosol in Wuhan Hospitals during COVID-19 Outbreak

**DOI:** 10.1101/2020.03.08.982637

**Authors:** Yuan Liu, Zhi Ning, Yu Chen, Ming Guo, Yingle Liu, Nirmal Kumar Gali, Li Sun, Yusen Duan, Jing Cai, Dane Westerdahl, Xinjin Liu, Kin-fai Ho, Haidong Kan, Qingyan Fu, Ke Lan

**Affiliations:** State Key Laboratory of Virology, Modern Virology Research Center, College of Life Sciences, Wuhan University, Wuhan, 430072, P. R. China; Division of Environment and Sustainability, The Hong Kong University of Science and Technology, Hong Kong SAR, P. R. China; Shanghai Environmental Monitoring Center, Shanghai 200235, P. R. China; School of Public Health, Key Lab of Public Health Safety of the Ministry of Education and Key Lab of Health Technology Assessment of the Ministry of Health, Fudan University, Shanghai 200032, P. R. China; JC School of Public Health and Primary Care, The Chinese University of Hong Kong, Hong Kong SAR, P. R. China

## Abstract

**Background:** The ongoing outbreak of COVID-19 has spread rapidly and sparked global concern. While the transmission of SARS-CoV-2 through human respiratory droplets and contact with infected persons is clear, the aerosol transmission of SARS-CoV-2 has been little studied.

**Methods:** Thirty-five aerosol samples of three different types (total suspended particle, size segregated and deposition aerosol) were collected in Patient Areas (PAA) and Medical Staff Areas (MSA) of Renmin Hospital of Wuhan University (Renmin) and Wuchang Fangcang Field Hospital (Fangcang), and Public Areas (PUA) in Wuhan, China during COVID-19 outbreak. A robust droplet digital polymerase chain reaction (ddPCR) method was employed to quantitate the viral SARS-CoV-2 RNA genome and determine aerosol RNA concentration.

**Results:** The ICU, CCU and general patient rooms inside Renmin, patient hall inside Fangcang had undetectable or low airborne SARS-CoV-2 concentration but deposition samples inside ICU and air sample in Fangcang patient toilet tested positive. The airborne SARS-CoV-2 in Fangcang MSA had bimodal distribution with higher concentration than those in Renmin during the outbreak but turned negative after patients number reduced and rigorous sanitization implemented. PUA had undetectable airborne SARS-CoV-2 concentration but obviously increased with accumulating crowd flow.

**Conclusions:** Room ventilation, open space, proper use and disinfection of toilet can effectively limit aerosol transmission of SARS-CoV-2. Gathering of crowds with asymptomatic carriers is a potential source of airborne SARS-CoV-2. The virus aerosol deposition on protective apparel or floor surface and their subsequent resuspension is a potential transmission pathway and effective sanitization is critical in minimizing aerosol transmission of SARS-CoV-2.

## Background

Circulating in China and 94 other countries and territories, the COVID-19 epidemic has resulted in 103,168 confirmed cases including 22,355 outside mainland China, with 3,507 deaths reported (March 7, 2020). Due to its increasing threat to global health, WHO has declared that the COVID-19 epidemic was a global public health emergency. The causative pathogen of the COVID-19 outbreak has been identified as a highly infectious novel coronavirus which is referred to as the Severe Acute Respiratory Syndrome Coronavirus 2 (SARS-CoV-2).^1–3^

The transmission of SARS-CoV-2 in humans is thought to be via at least 3 sources: 1) inhalation of liquid droplets produced by and/or 2) close contact with infected persons and 3) contact with surfaces contaminated with SARS-CoV-2.^4^ Moreover, aerosol transmission of pathogens has been shown in confined spaces.^5,6^ There are many respiratory diseases spread by the airborne route such as tuberculosis, measles and chickenpox.^7,8^ A retrospective cohort study conducted after the SARS epidemic in Hong Kong in 2003 suggested that airborne spread may have played an important role in the transmission of that disease.^9^ At present, there is little information on the characteristics of airborne SARS-CoV-2 containing aerosols, their concentration patterns and behaviour during airborne transmission due to the difficulties in sampling virus-laden aerosols and challenges in their quantification at low concentration. Such a lack of understanding limits effective risk assessment, prevention and control of COVID-19 disease outbreaks. This study on airborne SARS-CoV-2 was conducted in different areas inside two hospitals and public areas in Wuhan, China, the epicenter city during the initial disease outbreak. We aimed to 1) quantify the concentrations of airborne SARS-CoV-2 both inside the hospitals and in outdoor public areas, 2) evaluate the aerodynamic size distributions of SARS-CoV-2 aerosols that may mediate its airborne transmission, and 3) determine the dry deposition rate of the airborne SARS-CoV-2 in a patient ward room.

## Methods

### 1. Study design

This study is an experimental investigation on the concentration and aerodynamic characteristics of airborne SARS-CoV-2 aerosol in different areas of two hospitals: the Renmin Hospital of Wuhan University, designated for treatment of severe symptom COVID-19 patient during the disease outbreak and the Wuchang Fangcang Field Hospital, one of the first temporary hospitals which was renovated from an indoor sports stadium to quarantine and treat mildly symptom patients, and outdoor public areas in Wuhan during the coronavirus outbreak. We further classified the sampling locations into three categories according to their accessibility by different groups: 1) Patient Areas (PAA), where the COVID-19 patients have physical presence. These include the Intensive Care Units (ICU), Coronary Care Units (CCU) and ward rooms inside Renmin Hospital, a toilet and staff workstations inside Fangcang Hospital; 2) Medical Staff Areas (MSA), the workplaces in the two hospitals exclusively accessed by the medical staff who had direct contact with the patients and 3) Public Areas (PUA), which were venues open for the general public. The description and characteristics of sampling sites are shown in Table S1.

Three types of aerosol samples were collected: 1) Aerosol samples of total suspended particles (TSP) with no upper size limit to quantify RNA concentration of SARS-CoV-2 aerosol; 2) Aerodynamic size segregated aerosol samples to determine the size distribution of airborne SARS-CoV-2; 3) Aerosol deposition samples to determine the deposition rate of airborne SARS-CoV-2.

### 2. Sample collection

The sampling was conducted between February 17 and March 2, 2020 in the locations by two batches as shown in Table 1. All aerosol samples were collected on presterilized gelatin filters (Sartorius, Germany). Total of 30 TSP aerosol samples were collected on 25 mm diameter filters loaded into styrene filter cassettes (SKC Inc, US) and sampled air at a fixed flow rate of 5.0 litre per minute (LPM) using a portable pump (APEX2, Casella, US). Total of 3 size segregated aerosol samples were collected using a miniature cascade impactor (Sioutas impactor, SKC Inc., US) that separate aerosol into five ranges (> 2.5 μm, 1.0 to 2.5 μm, 0.50 to 1.0 μm and 0.25 to 0.50 μm on 25 mm filter substrates, and 0 to 0.25 μm on 37 mm filters) at a flow rate of 9.0 LPM. Total of 2 aerosol deposition samples were collected using 80 mm diameter filters packed into a holder with an effective deposition area of 43.0 cm^2^ and the filters were placed on the floor in two corners of Renmin Hospital ICU room intact for 7 days. Sampling durations and operation periods are detailed in Table S1. Prior to the field sampling, the integrity and robustness of experiment protocol was examined in the laboratory and described in Supplementary Appendix (Table S2).

**Table 1.**
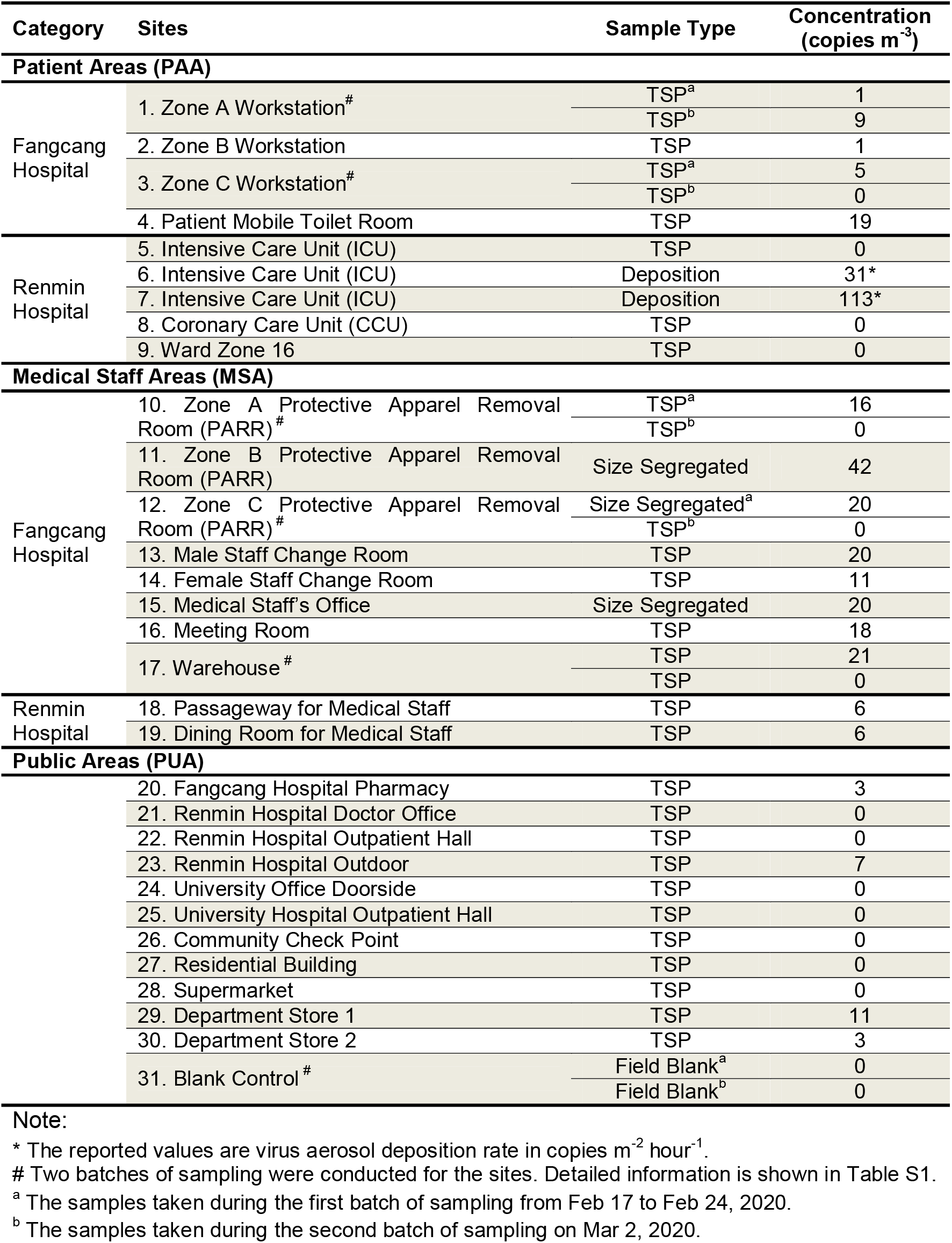
Concentration of airborne SARS-CoV-2 at different locations in Wuhan.

### 3. Analytical method and data analysis

After aerosol sample collection, all samples were handled immediately in the BSL-2 laboratory of Wuhan University. The 25, 37mm and 80 mm filter samples were dissolved in deionized water, then TRIzol LS Reagent (Invitrogen) was added to inactivate SARS-CoV-2 viruses and extract RNA according to the manufacturer’s instruction. First strand cDNA was synthesized using PrimeScript RT kit (TakaRa). Optimized ddPCR was used to detect the presence of SARS-CoV-2 viruses following our previous study.^10^ Analysis of the ddPCR data was performed with QuantaSoft software (Bio-Rad). The concentration reported by the procedure equals copies of template per microliter of the final 1x ddPCR reaction, which was normalized to copies m^−3^ in all the results, and hence the virus or viral RNA concentration in aerosol is expressed in copies m^−3^ hereafter. A detailed protocol is provided in Supplementary Appendix.

## Results

### 1. Airborne SARS-CoV-2 concentrations

The airborne SARS-CoV-2 concentrations in different categorized sites are shown in Table 1. The ICU, CCU and ward room in PAA of Renmin Hospital had negative test results. Fangcang Hospital workstations in different zones had low concentrations (1-9 copies m^−3^) of SARS-CoV-2 aerosol. The highest concentration in PAA of two hospitals was observed inside the patient mobile toilet room (19 copies m^−3^). In MSAs, the two sampling sites in Renmin Hospital had low concentration of 6 copies m^−3^, while the sites in Fangcang Hospital in general had higher concentrations. Particularly, the Protective Apparel Removal Rooms (PARRs) in three different zones inside Fangcang Hospital are among the upper range of airborne SARS-CoV-2 concentration from 18 to 42 copies m^−3^ in the first batch of sampling. During the second batch of sampling, the two TSP samples in the PARRs had negative test results with reduced number of medical staff and more rigorous sanitization processes in Fangcang. In PUA, SARS-CoV-2 aerosol concentrations were below 3 copies m^−3^, except for two occasions: one crowd gathering site near the entrance of a department store with frequent customer flow and one outdoor site next to Renmin Hospital with outpatients and passengers passing by.

### 2. Size distribution of SARS-CoV-2 aerosol

Figure 1 shows the SARS-CoV-2 aerosol concentrations in different aerodynamic size bins collected from PARRs in Zone B and C, and Medical Staff’s Office in Fangcang Hospital. The peak concentration of SARS-CoV-2 aerosols appears in two distinct size ranges, one in the submicron region with aerodynamic diameter dominant between 0.25 to 1.0 μm, and the other peak in supermicron region with diameter larger than 2.5 μm. The submicron region was dominantly noted in PARRs in Zone B and C of Fangcang Hospital (Figure 1a and 1b) with peak concentration of 40 and 9 copies m^−3^ in 0.25 to 0.5 μm and 0.5 to 1.0 μm, respectively. Whereas the supermicron region was observed in Fangcang Hospital Zone C PARR and Medical Staff’s Office (Figure 1b and 1c) with 7 and 9 copies m^−3^. The two concentration peaks in sub- and supermicron ranges have independent existence in SARS-CoV-2 aerosols and they do not necessarily co-exist indicating possible different formation mechanisms.

**Figure 1.**
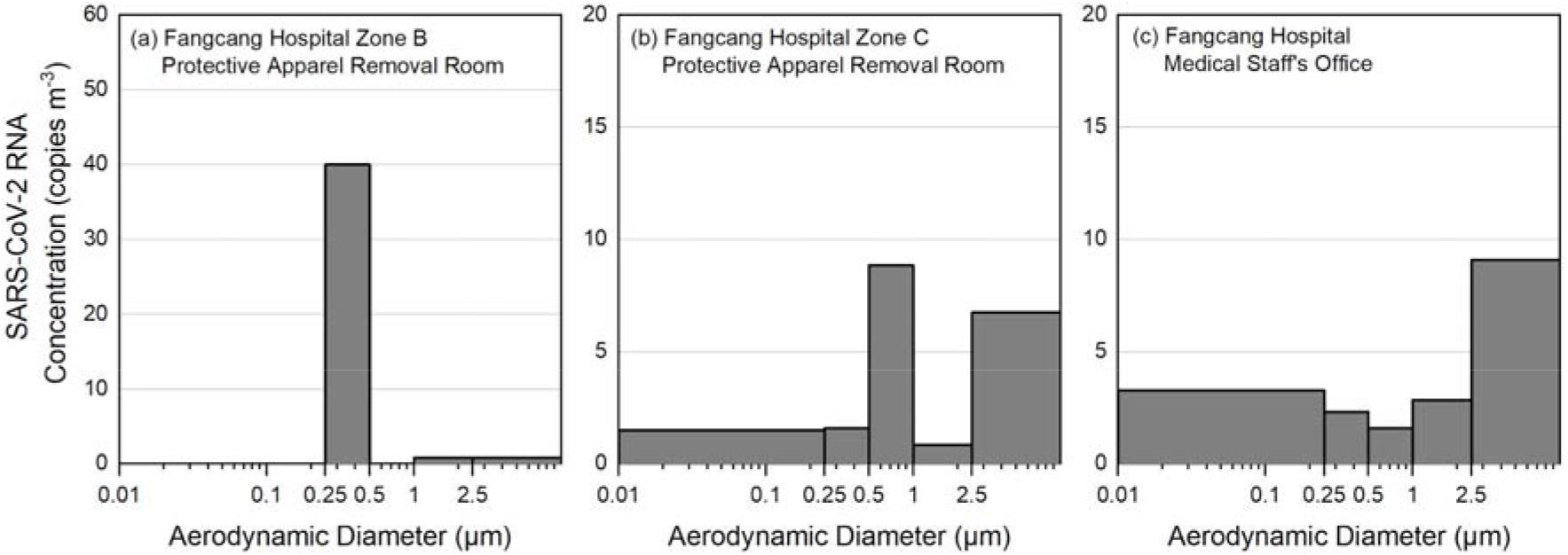
Concentration of airborne SARS-CoV-2 RNA in different aerosol size bins

### 3. Deposition rate of SARS-CoV-2 aerosol

The aerosol deposition sample collected from the Renmin Hospital ICU room had raw counts of SARS-CoV-2 RNA significantly above the detection limit as shown in Table S1, although the TSP aerosol sample concentration inside this ICU room was below detection limit during the 3 hour sampling period. The much longer integration time of 7 days for the deposition sample has contributed to the accumulation of virus sediment. The area normalized deposition rate inside the ICU room is calculated to be 31 and 113 copies m^−2^ hour^−1^. The sample with the higher deposition rate was placed in the hindrance-free corner of the room, approximately 3 meters from the patient’s bed. The other sample recorded lower virus copies and it was placed in another corner with medical equipment above, and approximately 2 meters from the patient’s bed. This may have blocked the path of virus aerosol sediment.

## Discussion

Generally undetectable or very low concentrations of airborne SARS-CoV-2 were found in most PAA inside the two hospitals in Wuhan. The negative pressure ventilation and high air exchange rate inside ICU, CCU and ward room of Renmin Hospital are effective in minimizing airborne SARS-CoV-2. Fangcang Hospital hosted over 200 mild symptom patients in each zone during the peak of the COVID-19 outbreak. However, the SARS-CoV-2 aerosol concentrations inside the patient hall were very low during the two batches of sampling periods, showing the protective and preventive measures taken in Fangcang Hospital are effective in hindering the aerosol transmission and reducing the potential infection risks of the medical staff. Inside the Renmin Hospital ICU rooms, the two aerosol deposition samples tested positive with an estimated deposition rate of 31 and 113 copies m^−2^ hr^−1^. The deposited virus may come from respiratory droplet or virus-laden aerosol transmission. Our findings add support to a hypothesis that virus-laden aerosol deposition may play a role in surface contamination and subsequent contact by susceptible people resulting in human infection.

This study also recorded an elevated airborne SARS-CoV-2 concentration inside the patient mobile toilet of Fangcang Hospital. This may come from either the patient's breath or the aerosolization of the virus-laden aerosol from patient’s faeces or urine during use. Ong et al. has found the wipe samples from room surfaces of toilets used by SARS-CoV-2 patients tested positive.^11^ Our finding has confirmed the aerosol transmission as an important pathway for surface contamination. We call for extra care and attention on the proper design, use and disinfection of the toilets in hospitals and in communities to minimize the potential source of the virus-laden aerosol.

MSAs in general have higher concentration of SARS-CoV-2 aerosol with biomodal size distributions compared to PAA in both hospitals during the first batch of sampling in the peak of COVID-19 outbreak. For Renmin Hospital sampling sites, the air circulation in MSA by design is isolated from that of the patient rooms. While for Fangcang Hospital, the non-ventilated temporary PARR has limited air penetration from the patient hall where the SARS-CoV-2 aerosol concentration was generally low. We believe one direct source of the high SARS-CoV-2 aerosol concentration may be the resuspension of virus-laden aerosol from the surface of medical staff protective apparel while they are being removed. These resuspended virus-laden aerosol originally may come from the direct deposition of respiratory droplets or virus-laden aerosol onto the protective apparel while medical staff having long working hours inside PAA, as shown from the SARS-CoV-2 deposition results in ICU room. Another possible source is the resuspension of floor dust aerosol containing virus that were transferred from PAA to MSA. The two virus-laden aerosol sources also appear to correspond to the sub- and supermicron peaks found in size-segregated samples. We hypothesize the submicron aerosol may come from the resuspension of virus-laden aerosol from staff apparel due to its higher mobility while the supermicron virus-laden aerosol may come from the resuspension of dust particles from the floors or other hard surfaces. The findings suggest virus-laden aerosols could first deposit on the surface of medical staff protective apparel and the floors in patient areas and are then resuspended by the movements of medical staff. The second batch of TSP samples taken in Fangcang MSAs all tested negative with reduced number of patients from > 200 to 100 per zone and implementation of more rigorous and thorough sanitization measures in Fangcang. The comparison of the two batches of samples showed the effectiveness and importance of sanitization in reducing the airborne SARS-CoV-2 in high risk areas.

In PUA outside the hospitals, we found the majority of the sites have undetectable or very low concentrations of SARS-CoV-2 aerosol, except for one crowd gathering site about 1 meter to the entrance of a department store with customers frequently passing through, and the other site next to Renmin Hospital where the outpatients and passengers passed by. It is possible that asymptomatic carriers of COVID-19 in the crowd may have contributed as the source of virus-laden aerosol during the sampling period.^12,13^ The results showed overall low risks in the public venues but do reinforce the importance of avoiding crowded gatherings and implementing early identification and diagnosis of asymptomatic carriers for early quarantine or treatment. Personal protection equipment such as wearing masks in public places or while in transit may reduce aerosol exposure and transmission.

The results from this study provide the first field report on the characteristics of airborne SARS-CoV-2 in Wuhan with important implications for the public health prevention and medical staff protection. We call for particular attentions on 1) the proper use and cleaning of toilets (e.g. ventilation and sterilization), as a potential spread source of coronavirus with relatively high risk caused by aerosolization of virus and contamination of surfaces after use; 2) for the general public, the proper use of personal protection measures, such as wearing masks and avoiding busy crowds; 3) the effective sanitization of the high risk area and the use of high level protection masks for medical staff with direct contact with the COVID-19 patients or with long stay in high risk area; 4) the renovation of large stadiums as field hospitals with nature ventilation and protective measures is an effective approach to quarantine and treat mild symptom patients so as to reduce the COVID-19 transmission among the public; 5) the virus may be resuspended from the contaminated protective apparel surface to the air while taking off and from the floor surface with the movement of medical staff. Thus, surface sanitization of the apparel before they are taken off may also help reduce the infection risk for medical staff.

## Acknowledgement

This study was supported by Special Fund for COVID-19 Research of Wuhan University. We are grateful to Taikang Insurance Group Co., Ltd, Beijing Taikang Yicai Foundation, Renmin Hospital and Wuchang Fangcang Hospital for their great support to this work. We would like to thank Prof. Hongmei Xu from Xi’an Jiaotong University, Qingdao Laoying Environmental Technology Co., Ltd, Beijing Top Science Co.,Ltd, Shanghai Leon Scientific Instrument Co.,Ltd, Shanghai Eureka Environmental Protection Hi-tech. Ltd, Sapiens Environmental Technology Co., Ltd for their support in providing the sampling devices and technical support in this study. The authors also thank Cuiping Wang, Qingli Zhang, Guoping Liang, Zhao Song for their assistance in filter sample preparation and logistics support.

