## Supplementary Appendix for "Aerodynamic Characteristics and RNA Concentration of SARS-CoV-2 Aerosol in Wuhan Hospitals during COVID-19 Outbreak"

### Methods

#### 1. Sample collection

All aerosol samples were collected by presterilized gelatin filters with pore size 3 μm (Sartorius, Germany) for two considerations. First, the gelatin filters have demonstrated high collection efficiency for virus aerosol without strong relevance to the aerodynamic size dependent particle inertial collection, which is important when no prior knowledge on the size of airborne virus aerosol is available.^1^ Second, the gelatin filters have high recovery efficiency with direct dissolution into ultra-small volume of water to enhance the sample concentration for RNA quantification.^2^

The TSP aerosol samples were collected on 25 mm diameter gelatin filters loaded in clear styrene filter cassette (SKC Inc, US). The air sample was drawn through the filter at a fixed flow rate of 5.0 litre per minute (LPM) using Casella portable pump (APEX2, Casella, US). The aerodynamic size segregated aerosol samples were collected using a miniature cascade impactor (Sioutas impactor, SKC Inc., US) loaded with four impaction stages that separate the particle aerodynamic size into five size ranges (> 2.5 μm, 1.0 to 2.5 μm, 0.50 to 1.0 μm and 0.25 to 0.50 μm under impaction stage, and 0- 0.25 μm as filtration stage) at a flow rate of 9.0 LPM. 25 mm and 37 mm gelatin filters were used for four impaction stages and one filtration stage, respectively. The effective air velocity through the 25 mm gelatin filter for TSP aerosol sample and 37 mm gelatin filter for cascade filtration stage are 0.22 and 0.18 meter/second, respectively, well within the specification of gelatin filter for high efficiency virus collection while maintaining the filter integrity. The flow rates of all the samplers were checked with Drycal flow meter (Defender 510, Mesa Labs, US) and adjusted to nominal flow within ± 5% prior to the sampling. To prevent sample contamination, all the filters were preloaded inside the samplers in Class 100 sterilized room and sealed with Teflon tapes. Laboratory and field blanks were prepared following the identical procedures. The aerosol deposition sample was collected using an 80 mm diameter gelatin filter packed in a holder with an effective deposition area of 43.0 cm^2^.

#### 2. Analytical Methods

**2.1 Aerosolization and sampling of test EV71 virus particles in the laboratory**

Prior to the field sampling for the SARS-CoV-2 aerosol samples, the integrity and robustness of experiment protocol was examined in the laboratory for the virus aerosol collection by the gelatin filters and the subsequent filter processing using EV71 virus as a surrogate. First, 20µl of EV71 viruses stock (3×10^7^ TCID_50_) was diluted in 2 ml deionized water, then pipetted into a sterilized glass vessel and aerosolized by a nebulizer (Yuwell, China) in BSL-2 cabinet. After 10 minutes sedimentation of the aerosol, the TSP aerosol sampling device was set up with air inlet at 1 m distance and the same height of the nebulizer. The sampler operated at a flow rate of 5.0 LPM with 25 mm diameter gelatin filter following the identical protocol as in this field study. Each sampling lasted for 1 h and three independent samplings were conducted. Gelatin filters processing, RNA extraction and cDNA synthesis procedures were the same as other air samples. 1µl pure EV71 virus stock was used as positive control for subsequent analysis. Real-time PCR (RT-PCR) system (Applied Biosystems) was used to quantify the viral load in EV71 air samples. Thermal cycling was performed as follows: 95°C for 10 min and then 40 cycles of 95 °C for 15 s, 60 °C for 30 s. Table S2 shows the results comparison between the positive control and air samples with consistent results proving the integrity of the designed sampling and processing protocol.

**2.2 RNA extraction, cDNA synthesis and primer selection**

For gelatin filters with the collected air samples, each gelatin was transferred to a clean 1.5 ml tube (15 ml conical tube for 80 mm gelatin filters) right after the sampling campaign and 400 µl (4 ml for 80 mm gelatin filters) sterile deionized water per sample was added immediately to each tube. Then the tubes were centrifuged and incubated at 37°C for 10 minutes by a block heater to dissolve the gelatin. All samples were inactivated using a 4:1 ratio of TRIzol LS Reagent (Invitrogen), total RNA was extracted according to the manufacturer’s instruction. RNA was dissolved in 30 µl RNase-free DI water per sample. First strand cDNA was synthesized using PrimeScript RT Master Mix (TakaRa) with random primer and oligo dT primer. In accordance with current clinical diagnosis of COVID-19 in China, primers and probes that targeted the ORF1ab and N genes of SARS-CoV-2 were used according to Chinese Center for Disease Control and Prevention (CCDC), and the sum of ORF1ab and N primer/probe sets results was considered as a representation of virus.

The primers and probes selected were targeted at the ‘ORF1ab’ and ‘N’ genes of SARS-CoV-2.^3^

ORF1ab:	forward primer:5'-CCCTGTGGGTTTTACACTTAA-3'

reverse primer:5'-ACGATTGTGCATCAGCTGA-3'

Probe: 5'-FAM-CCGTCTGCGGTATGTGGAAAGGTTATGG-BHQ1-3'

N:	forward primer: 5'-GGGGAACTTCTCCTGCTAGAAT-3'

reverse primer: 5'-CAGACATTTTGCTCTCAAGCTG-3'

Probe: 5'-FAM-TTGCTGCTGCTTGACAGATT-TAMRA-3'

**2.3 Droplet Digital Polymerase Chain Reaction (ddPCR)**

The ddPCR was performed according to the manufacturer’s instructions for the QX200 Droplet Digital PCR System (Bio-Rad) using supermix for probe (without dUTP) (Bio-Rad).^4,5^ Briefly, the TaqMan PCR reaction mixture was made from a 2x supermix for probe (without dUTP), 20x primer and probes (final concentrations of 900 and 250 nM, respectively) and different volumes of template in a final volume of 20 μl. It was then converted to droplets with the QX200 droplet generator, and transferred to a 96-well plate, sealed and cycled in a T100 Thermal Cycler (Bio-Rad) using cycling protocol: 95 °C for 10 min, followed by 40 cycles of 94 °C for 30 s (denaturation) and 60 °C for 1 min (annealing) followed by an infinite 4-degree hold. The cycled plate was then automatically read in the FAM channels using the QX200 reader.

Analysis of the ddPCR data was performed with QuantaSoft analysis software v.1.7.4.0917 (Bio-Rad) that accompanied the droplet reader to calculate the concentration of the target sequences, along with their Poisson-based 95 % confidence intervals. The positive populations for each primer/probe are identified using positive and negative controls with single (i.e., not multiplexed) primer–probe sets. The lower limit of detection (LLoD) of the optimized ddPCR is 2.18 copies and 0.42 copies per reaction (20 µl) for ORF1ab and N primers/probe sets, respectively.^6^

**Table S1. Specifications of sampling sites**

| **Category** | **Sites** | **Ventilation Type** | **Area Size#** | **Inpatients number/ symptom*** | **Sampling Period** |
| --- | --- | --- | --- | --- | --- |
| **Patient Areas (PAA)** | |  |  |  |  |
| Fangcang Hospital | 1. Zone A Workstation | Indoor; Nature Ventilation | > 500m^2^ | >200/ Mild | 20:00 02/23 - 12:00 02/24^a^ |
|  |  |  |  | <100/ Mild | 9:00 03/02 - 20:00 03/02^b^ |
|  | 2. Zone B Workstation | Indoor; Nature Ventilation | > 500m^2^ | >200/ Mild | 20:00 02/23 - 12:00 02/24 |
|  | 3. Zone C Workstation | Indoor; Nature Ventilation | > 500m^2^ | >200/ Mild | 20:00 02/23 - 12:00 02/24^a^ |
|  |  |  |  | <100/ Mild | 9:00 03/02 - 20:00 03/02^b^ |
|  | 4. Patient Mobile Toilet Room | Enclosed Room; No Ventilation | ~1 m^2^ |  | 20:00 02/23 - 12:00 02/24 |
| Renmin Hospital | 5. Intensive Care Unit (ICU) | Indoor; Negative Pressure | 16 m^2^ | 1/ Severe | 9:30 02/18 - 12:30 02/18 |
|  | 6. Intensive Care Unit (ICU) | Indoor; Negative Pressure | 16 m^2^ | 1/ Severe | 10:00 02/18 - 11:45 02/25 |
|  | 7. Intensive Care Unit (ICU) | Indoor; Negative Pressure | 16 m^2^ | 1/ Severe | 10:00 02/18 - 11:45 02/25 |
|  | 8. Coronary Care Unit (CCU) | Indoor; Negative Pressure | 16 m^2^ | 1/ Severe | 9:30 02/18 - 12:30 02/18 |
|  | 9. Ward Zone 16 | Indoor; Negative Pressure | 16 m^2^ | 2/ Severe | 9:30 02/18 - 12:30 02/18 |
| **Medical Staff Areas (MSA)** | |  |  |  |  |
| Fangcang Hospital | 10. Zone A Protective Apparel Removal Room (PARR) | Indoor; Small Air Purifier | 5 m^2^ |  | 20:00 02/23 - 12:00 02/24^a^ |
|  |  |  |  |  | 9:00 03/02 - 20:00 03/02^b^ |
|  | 11. Zone B Protective Apparel Removal Room (PARR) | Indoor; Small Air Purifier | 5 m^2^ |  | 20:00 02/23 - 12:00 02/24 |
|  | 12. Zone C Protective Apparel Removal Room (PARR) | Indoor; Small Air Purifier | 5 m^2^ |  | 20:00 02/23 - 12:00 02/24^a^ |
|  |  |  |  |  | 9:00 03/02 - 20:00 03/02^b^ |
|  | 13. Male Staff Change Room | Indoor; Nature Ventilation | 150 m^2^ |  | 17:30 02/23 - 20:30 02/23 |
|  | 14. Female Staff Change Room | Indoor; Nature Ventilation | 150 m^2^ |  | 17:30 02/23 - 20:30 02/23 |
|  | 15. Medical Staff’s Office | Indoor; Nature Ventilation | 12 m^2^ |  | 20:00 02/23 - 12:00 02/24 |
|  | 16. Meeting Room | Indoor; Mechanical Ventilation | 200 m^2^ |  | 17:30 02/23 - 20:30 02/23 |
|  | 17. Warehouse | Indoor; Nature Ventilation | 25 m^2^ |  | 17:30 02/23 - 20:30 02/23^a^ |
|  |  |  |  |  | 9:00 03/02 - 20:00 03/02^b^ |
| Renmin Hospital | 18. Passageway for Medical Staff | Indoor | 10 m^2^ |  | 9:30 02/18 - 11:00 02/18 |
|  | 19. Dining Room for Medical Staff | Indoor | 20 m^2^ |  | 9:30 02/18 - 11:00 02/18 |
| **Public Areas (PUA)** | |  |  |  |  |
|  | 20. Fangcang Hospital Pharmacy | Indoor; Mechanical Ventilation | 25 m^2^ |  | 17:30 02/23 - 20:30 02/23 |
|  | 21. Renmin Hospital Doctors’ Office | Outdoor; Nature Ventilation | 20 m^2^ |  | 9:30 02/18 - 11:00 02/18 |
|  | 22. Renmin Hospital Outpatient Hall | Outdoor; Nature Ventilation | 800 m^2^ |  | 9:30 02/18 - 12:30 02/18 |
|  | 23. Renmin Hospital Outdoor | Outdoor |  |  | 9:30 02/18 - 11:00 02/18 |
|  | 24. University Office Doorside | Indoor; Nature Ventilation | 20 m^2^ |  | 15:00 02/17 - 16:30 02/17 |
|  | 25. University Hospital Outpatient Hall | Indoor; Nature Ventilation | 30 m^2^ |  | 12:00 02/19 - 18:00 02/19 |
|  | 26. Community Check Point | Outdoor |  |  | 12:00 02/19 - 18:00 02/19 |
|  | 27. Residential Building | Outdoor |  |  | 12:00 02/19 - 18:00 02/19 |
|  | 28. Supermarket | Outdoor |  |  | 18:00 02/17 - 19:30 02/17 |
|  | 29. Department Store 1 | Outdoor |  |  | 18:00 02/17 - 19:30 02/17 |
|  | 30. Department Store 2 | Outdoor |  |  | 12:00 02/19 - 18:00 02/19 |
|  | 31. Blank Control |  |  |  | 18:00 02/17^a^ |
|  |  |  |  |  | 9:00 03/02^b^ |

\# The height of Workstations in Zone A and Zone C of Fangcang Hospital is about 10 m; the height of Workstation in Zone B of Fangcang Hospital is about 4 m. Heights of other sampling sites are around 2.5 -3 m.

* Inpatients number and symptom severity on the date of sampling.

^a^ The samples taken during the first batch of sampling from Feb 17 to Feb 24, 2020.

^b^ The samples taken during the second batch of sampling on Mar 2, 2020.

**Table S2. Cycle threshold value (Ct) of EV71 virus positive control and laboratory test air samples RT-PCR.** The Ct or threshold cycle value is the cycle number at which the fluorescence generated within a reaction crosses the threshold in real-time PCR. Lower Ct values correspond to higher viral loads.

| **Replication** | **1µl positive control** | **Air samples** |
| --- | --- | --- |
| 1 | 14.35 | 21.23 |
| 2 | 13.93 | 21.13 |
| 3 | 13.75 | 21.13 |
